# Tau phosphorylated at serine 356 is associated with Alzheimer’s disease pathology and can be lowered in mouse and human brain tissue using the NUAK inhibitor WZ4003

**DOI:** 10.1101/2023.08.28.553851

**Authors:** Lewis W. Taylor, Elizabeth M. Simzer, Claire Pimblett, Oscar T.T. Lacey-Solymar, Robert I. McGeachan, Soraya Meftah, Jamie L. Rose, Maxwell P. Spires-Jones, James H. Catterson, Henner Koch, Imran Liaquat, Jonathan H. Clarke, John Skidmore, Sam A. Booker, Paul M. Brennan, Tara L. Spires-Jones, Claire S. Durrant

## Abstract

Tau hyperphosphorylation and aggregation is a common feature of many dementia-causing neurodegenerative diseases. Tau can be phosphorylated at up to 85 different sites, and there is increasing interest in whether tau phosphorylation at specific epitopes, by specific kinases, plays an important role in disease progression. The AMP-activated protein kinase (AMPK) related enzyme NUAK1 been identified as a potential mediator of tau pathology, whereby NUAK1-mediated phosphorylation of tau at Ser356 prevents the degradation of tau by the proteasome, further exacerbating tau hyperphosphorylation and accumulation. This study provides a detailed characterisation of the association of p-tau Ser356 with progression of Alzheimer’s disease pathology, identifying a Braak stage-dependent increase in p-tau Ser356 protein levels and an almost ubiquitous presence in neurofibrillary tangles. We also demonstrate, using sub-diffraction-limit resolution array tomography imaging, that p-tau Ser356 co-localises with synapses in AD post-mortem brain tissue, increasing evidence that this form of tau may play important roles in AD progression. To assess the potential impacts of pharmacological NUAK inhibition in an *ex vivo* system that retains multiple cell types and brain-relevant neuronal architecture, we treated postnatal mouse organotypic brain slice cultures from wildtype or APP/PS1 littermates with the commercially available NUAK1/2 inhibitor WZ4003. Whilst there were no genotype specific effects, we found that WZ4003 results in a culture-phase dependent loss of total tau and p-tau Ser356, which corresponds with a reduction in neuronal and synaptic proteins. By contrast, application of WZ4003 to live human brain slice cultures results in a specific lowering of p-tau Ser356, alongside increased neuronal tubulin protein. This work identifies differential responses of postnatal mouse organotypic brain slice cultures and adult human brain slice cultures to NUAK1 inhibition that will be important to consider in future work developing tau-targeting therapeutics for human disease.

## Introduction

Hyperphosphorylation and aggregation of tau is a key feature of a number of dementia-causing diseases, including primary tauopathies, such as Frontotemporal dementias (FTD) with tau pathology, and secondary tauopathies, such as Alzheimer’s disease (AD), where accumulation of amyloid-beta (Aβ) is thought to initiate the disease cascade^1,2^. Whilst tau plays a number of important physiological roles in the brain, including regulating microtubule function, myelination, neuronal excitability and DNA protection^3^, there is strong evidence that hyperphosphorylated, oligomeric tau disrupts synaptic function and may be an important driver of synapse loss and neurodegeneration in dementia^4,5^. Tau can be phosphorylated at up to 85 different sites (45 serine, 35 threonine and 5 tyrosine residues), with the levels of phosphorylation regulated by an ever expanding list of kinases and phosphatases^6–8^. There is increasing focus on whether specific forms of phosphorylated tau are key drivers of downstream pathology, and whether targeting upstream kinases could be an effective therapeutic tool to mitigate tau pathology in dementia^6,9^.

Of recent interest is the possibility that increased levels of the AMP-activated protein kinase (AMPK) related kinase, NUAK1, in AD and primary tauopathies, results in the specific phosphorylation of tau at Ser356^9^. Ser356 is located in repeat 4 of the microtubule binding domains, so phosphorylation of this site is likely to disrupt key aspects of tau function^10^. Interestingly a mutation in this site on the *MAPT* gene has been linked to a very-early onset form of FTD with Parkinsonism linked to chromosome 17 (FTDP-17)^11–13^ and studies in Drosophila have highlighted that p-tau Ser356 could be a catalyst for further downstream phosphorylation and aggregation^14,15^. NUAK1-mediated phosphorylation of tau at Ser356 has also been identified as a mechanism for regulating total tau levels, with the chaperone C-terminus of Hsc70-interacting protein (CHIP) unable to bind tau phosphorylated at Ser356, thus preventing ubiquitination of tau and its subsequent degradation by the proteasome^9,16,17^. By crossing NUAK1^+/-^ mice to the tauopathy P301S mouse model, Lasagna-Reeves et al. found that reduction of NUAK1 lowered both p-tau Ser356 and total tau levels and rescued aspects of tau pathology^9^. This work highlighted NUAK1 as an attractive target for therapeutic development in primary tauopathies, opening important questions about whether similar strategies could be applicable to secondary tauopathies such as AD.

Whilst there are reports that tau is phosphorylated at Ser356 in end-stage AD^10,18–21^, the progression of p-tau Ser356 accumulation over the disease time course, its representation in tangles and its association with synapses (synaptic tau has been found to be important for both tau toxicity and trans-synaptic tau spread^22–25)^ has not been fully characterised. Of the few studies that do examine appearance of this epitope in AD brain tissue, many use the 12E8 antibody^26,27^, which shows considerable preference for p-tau Ser262 complicating interpretation of the unique involvement of p-tau Ser356^28^. In this work, we characterise specific accumulation of p-tau Ser356 in AD brain using biochemical and histological methods including sub-diffraction limit-resolution microscopy to examine synapses^22,29^.

Based on the likely pathological involvement of p-tau Ser356, we further explore the effects of pharmacological inhibition of phosphorylation of tau at this residue. We characterise the impact of the commercially available NUAK inhibitor WZ4003, which has been previously shown to inhibit NUAK1 activity *in vitro*^30^ and reduce p-tau Ser356 in neuroblastoma cells^9^. In this study, we look to examine the impact of WZ4003 treatment under a number of physiological and pathological conditions using both mouse and human organotypic brain slice cultures, which retain physiologically relevant neuronal architecture, supporting cell types and synaptic connections for several weeks *in vitro*^31–35^. Our results reinforce the importance of p-tau Ser356 in AD and highlight potential biological differences in mouse and human brain in terms of how NUAK1 regulates tau turnover (experiment overview in **Fig. 1**).

**Figure 1:**
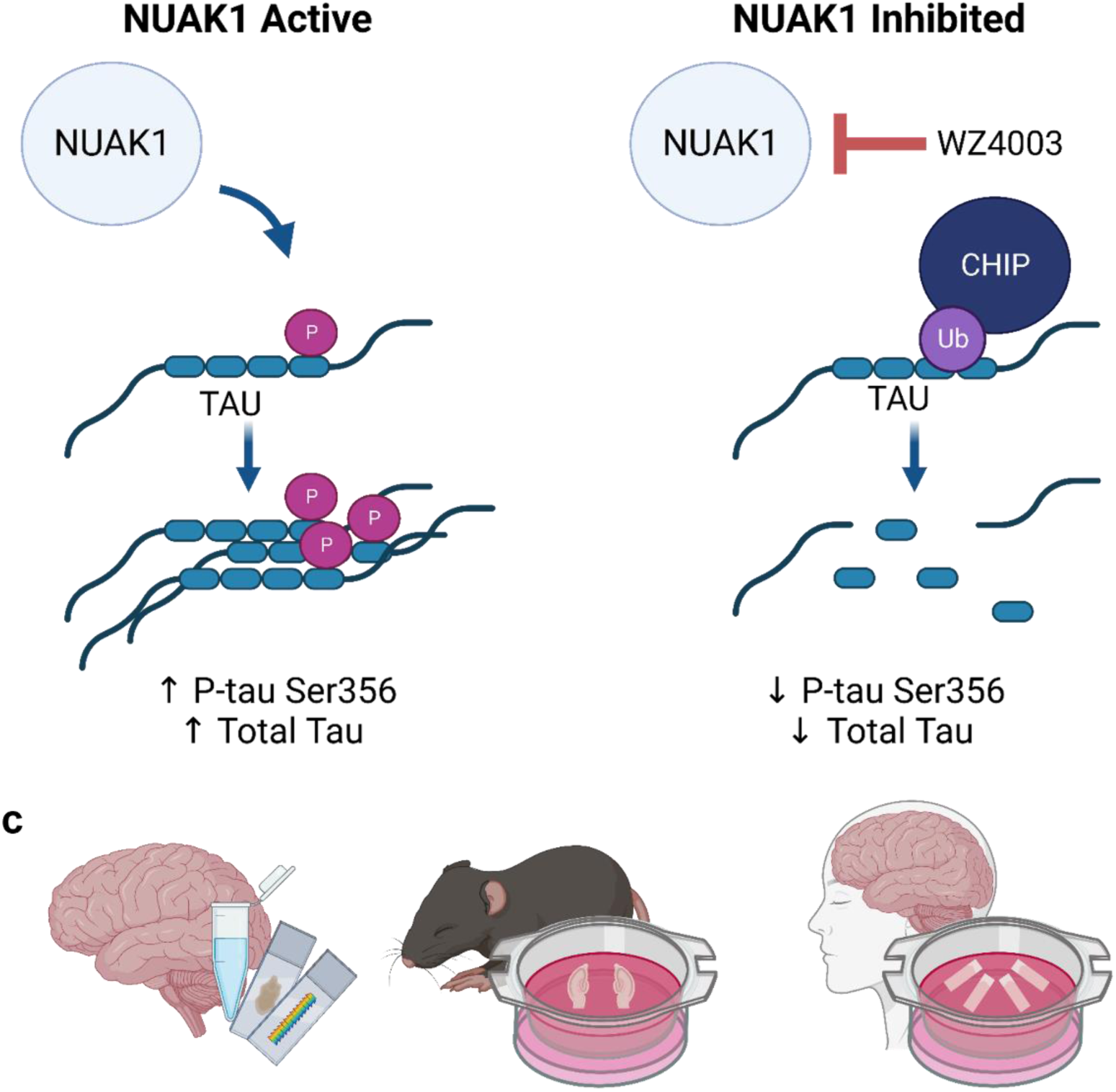
Proposed mechanism by which NUAK1 activity regulates tau levels via phosphorylation at Ser356. ***(a)** Under cases where NUAK1 is active (or overactive as proposed in AD), NUAK1 phosphorylates tau at Ser356, located in the 4^th^ microtubule binding repeat. CHIP is unable to bind tau phosphorylated at Ser356, allowing p-tau Ser356 and ultimately total tau to accumulate. **(b)** When NUAK1 is inhibited (such as with WZ4003), there is less phosphorylation at Ser356. CHIP is able to bind to tau and targets it for degradation by the proteosome. **(c)** This work uses multiple experimental systems, including post-mortem human brain (assessed by Western blot, paraffin section staining and array tomography), mouse organotypic brain slice cultures (MOBSCs) and human brain slice cultures (HBSCs). Cartoons generated using BioRender*

## Materials and Methods

### Human post-mortem brain tissue

All post-mortem brain tissue used in this study was obtained with ethical approval from the Edinburgh Sudden Death Brain Bank. This study was reviewed and approved by the Edinburgh Brain Bank ethics committee, a joint office for NHS Lothian and the University of Edinburgh, under ethical approval number 15-HV-016. The Edinburgh Brain Bank is a Medical Research Council funded facility with research ethics committee (REC) approval (16/ES/0084). Individual demographics for human post-mortem studies are listed in **Table 1** and case numbers/ types of tissue are detailed in methods section below for each experiment. Inclusion criteria for AD cases were: clinical dementia diagnosis, Braak stage 5-6 and a post-mortem neuropathological diagnosis of AD. Control subjects were chosen to be age/ sex matched as much as possible, with inclusion criteria of no diagnosed neurological or psychiatric condition. Exclusion criteria for both AD and control samples were substantial non-AD neuropathological findings (such as dementia with Lewy bodies/ haemorrhage).

**Table 1:**
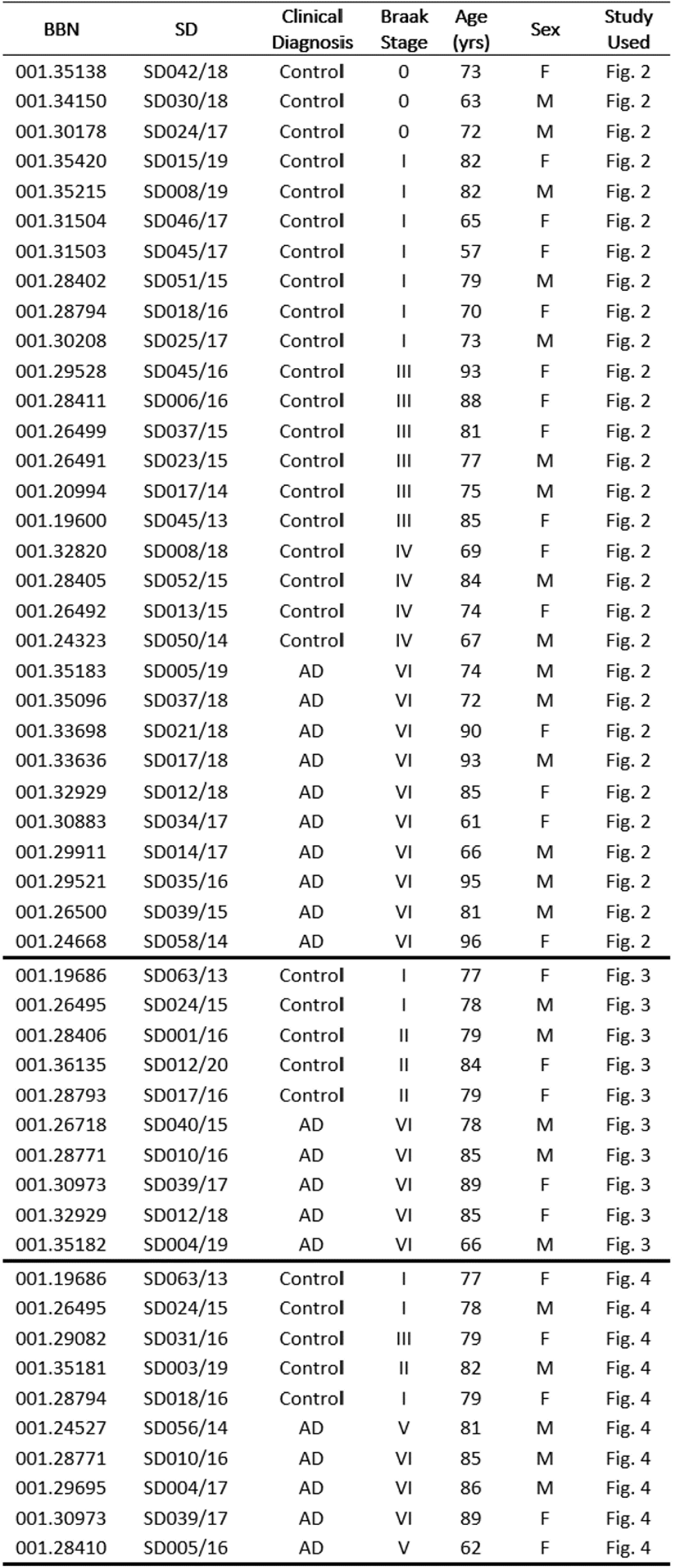
Demographic and neuropathological characteristics of all human post-mortem subjects. AD= Alzheimer’s disease, BBN= Medical Research Council Brain Bank Number, SD= Edinburgh Brain Bank Number.

### Human post-mortem total protein homogenate and synaptoneurosome preparation

Ten different frozen post-mortem human brain case samples were obtained per Braak Stage for the BA20/21 region (summary details listed in **Table 2)**. Total protein homogenate and synaptoneurosomes were generated as previously described^36^. For each case sample, 300-500mg of frozen tissue was thawed and immediately homogenised, on ice, in a glass-teflon dounce homogeniser in 1ml of homogenisation buffer (25 mM HEPES (pH 7.5), 120 mM NaCl, 5 mM KCl, 1 mM MgCl2, 2 mM CaCl2, protease and phosphatase inhibitors (Roche complete mini: 11836153001)). The homogenate was then flushed through an 80μm-pore nylon filter (Millipore NY8002500), with 300μl of the resulting crude total homogenate set aside and frozen at -80°C. To generate synaptoneurosome preps, total homogenate was further filtered through a 5μm filter (Millipore SLSV025NB). Synaptoneurosomes were centrifuged at 1000xg for 5 minutes and the pellet collected. Proteins (either total homogenate or synaptoneurosome) were extracted in an SDS buffer (100mM Tris-HCl, 4% SDS) and the protein concentration determined by commercial BCA assay before equal protein amounts were diluted in 2x Laemmli buffer, boiled then loaded for Western blot (below).

**Table 2:** Summary demographic information of human post-mortem subjects (homogenate Western blot study (Figure 2)). Cartoons generated using BioRender.

| Braak Stage: | 0-I<br>(N=10) | III-IV<br>(N=10) | VI<br>(N=10) | Overall<br>(N=30) |
| --- | --- | --- | --- | --- |
| **Age (Years)** |  |  |  |  |
| Mean (SD) | 71.6 (8.22) | 79.3 (8.37) | 81.3 (12.5) | 77.4 (10.5) |
| Median [Min, Max] | 72.5 [57.0, 82.0] | 79.0 [67.0, 93.0] | 83.0 [61.0, 96.0] | 76.0 [57.0, 96.0] |
| **Sex** |  |  |  |  |
| F | 5 (50.0%) | 6 (60.0%) | 4 (40.0%) | 15 (50.0%) |
| M | 5 (50.0%) | 4 (40.0%) | 6 (60.0%) | 15 (50.0%) |

### Human post-mortem immunohistochemistry

**(Summary patient details in Table 3)** 4 μm sections of formalin-fixed, paraffin-embedded tissue from Broadmann area 20/21 (inferior temporal gyrus) were de-waxed in xylene then decreasing concentrations of ethanol, incubated with autofluorescence inhibitor reagent (Millipore 2160), and non-specific antigens blocked by incubation in a solution of 0.1 M PBS containing 0.3% Triton X-100 and 10% normal donkey serum for 1 hour. Primary antibodies to p-tau Ser356 (Abcam: ab75603, diluted 1:1000) and GFAP (Abcam: ab4674, diluted 1:3000) were diluted in blocking buffer and incubated on sections overnight at 4 °C. Sections were washed with 0.1 M PBS containing 0.3% triton X-100 then incubated in secondary antibodies (donkey anti-rabbit Alexa-fluor 594, Invitrogen A21207 and donkey anti chicken Alexa-fluor 647, Invitrogen A78952) diluted 1:200 in block buffer at room temperature for 1 hour. Sections were washed with PBS then incubated in 0.5% Thioflavin S in 50% ethanol, 50% H2O solution for 8 minutes in the dark at room temperature. Slides were dipped in 80% ethanol to reduce Thioflavin S background then rinsed thoroughly in distilled water before coverslipping with Immu-Mount^TM^ (Epredia^TM^: 9990402). Sections were imaged with a 20x 0.8NA objective on a Zeiss AxioImager Z2 microscope equipped with a CoolSnap camera. Ten regions of interest were imaged in the cortex of each slide sampling in a systematic random fashion to sample throughout all 6 cortical layers. Images were examined in Fiji (ImageJ) by a blinded experimenter and the presence or absence of p-tau Ser536 noted in each neurofibrillary tangle identified by thioflavin S staining in classic “flame” shape in the size expected for neuronal tangles.

### Human post-mortem array tomography

Tissue from five non-AD control patients, with Braak Stages I-II and Thal phase 1-2, and five AD patients with Braak Stage VI and Thal phase 5 were acquired (summary patient details in **Table 4**). As described previously^22^ fresh tissue was fixed in 4% paraformaldehyde for 3 hours, dehydrated in ethanol and embedded in LR White Resin. A diamond knife (Diatome) mounted onto an Ultracut microtome (Leica) was used to cut the embedded tissue into 70nm serial sections. 15-30 serial section ribbons were collected onto gelatin-coated coverslips and immunostained with the following primary antibodies for 1 hour: 1:500 Rabbit p-tau Ser356 (Abcam: ab75603), 1:100 goat synaptophysin (R&D Systems: AF555), and 1:100 mouse AT8 (ThermoFisher: MN1020). Secondary antibodies, donkey anti-rabbit Alexa-fluor 405 (Abcam: ab175651), donkey anti-goat Alexa-fluor 594 (Abcam: ab150136), and donkey anti-mouse Alexa-fluor 647 (ThermoFisher: A32787) were then applied at 1:50 concentration to ribbons for 45 minutes. Images were obtained with a 63x 1.4 NA objective on a Leica TCS confocal microscope. Images from the same locus in each serial section along a ribbon were then aligned, thresholded, and parameters quantified using in-house scripts in Fiji (ImageJ) and MATLAB. Resulting parameter data was statistically analysed using custom R Studio scripts. Analysis code is available on GitHub (https://github.com/Spires-Jones-Lab). Imaris software was used to generate 3-D reconstructions of serial section co-localisations.

### Animals

APP/PS1 (APPswe, PSEN1dE9) and wildtype litter mate male and female mouse pups, aged 6-9 days old (P6-9), were obtained from a breeding colony at the Bioresearch and Veterinary Services Animal Facility at the University of Edinburgh. Animals were culled via cervical dislocation, performed by a trained individual, who was assessed by the Named Training and Competency Officer (NTCO). All animal work was conducted according to the Animals (Scientific Procedures) Act 1986 under the project licence PCB113BFD and PP8710936. All animals were bred and maintained under standard housing conditions with a 12/12 hour light-dark cycle.

### Mouse organotypic brain slice culture generation and maintenance

Mouse organotypic brain slice cultures (MOBSCs) were generated and maintained as described previously^31–34^, with minor modifications. Mouse pups aged postnatal day (P)6-9 were culled by cervical dislocation. Brains were rapidly transferred to ice-cold 0.22 μm-filtered dissection medium composed of 87 mM NaCl, 2.5 mM KCl, 25 mM NaHCO3, 1.25 mM NaH2PO4, 25 mM glucose, 75 mM sucrose, 7 mM MgCl2, 0.5 mM CaCl2, 1 mM Na-Pyruvate, 1 mM Na-Ascorbate, 1 mM kynurenic acid and 1X penicillin/streptomycin (ThermoFisher: 15140122) (340 mOsm, pH 7.4), bubbled with 95% O2, 5% CO2. All salts and chemicals were purchased from Merck. Brains were then mounted on and glued (cyanoacrylate, Loctite) to a vibratome stage. A Leica VT1200S vibratome was used to cut 350 µm-thick horizontal slices, from which the hippocampus was dissected with fine needles. Hippocampal slices were plated on membranes (Millipore: PICM0RG50) sitting on top of 1 ml of maintenance medium, placed inside 35 mm culture dishes. The maintenance medium was 0.22 µm-filtered and composed of MEM with Glutamax-1 (50%) (Invitrogen: 42360032), heat-inactivated horse serum (25%) (ThermoFisher: 26050070), EBSS (18%) (ThermoFisher: 24010043), D-glucose (5%) (Sigma: G8270), 1X penicillin/streptomycin (ThermoFisher: 15140122), nystatin (3 units/ml) (Merck: N1638) and ascorbic acid (500 μM) (Sigma-Aldrich: A4034). Slice cultures were then immediately placed into an incubator and maintained at 37°C with 5% CO2 thereafter. The slice culture medium was changed fully within 24 hours of plating, then at 4 days in vitro (*div*), 7 *div* and weekly thereafter. 4 culture dishes were made per pup, with 1- 2 slices plated per dish. Cultures were kept for either 2 weeks or 4 weeks, with 5 μM WZ4003 (APExBIO: B1374) or DMSO (Sigma-Aldrich: D2438) control applied in culture medium at every feed during the treatment period (either 0-2 weeks or 2-4 weeks).

### Human brain slice cultures

Human brain slice cultures (HBSCs) were generated from surplus neocortical access tissue from patients undergoing tumour resection surgery with ethical approval from the Lothian NRS Bioresource (REC number: 15/ES/0094, IRAS number: 165488) under approval number SR1319. Additional approval was obtained for receiving data on patient sex, age, reason for surgery and brain region provided (NHS Lothian Caldicott Guardian Approval Number: CRD19080). Patient details are listed in **Table 5**. The informed consent of patients was obtained using the Lothian NRS Bioresource Consent Form. Dissection and culture methods have been adapted from published studies^37–40^. Access, non-tumour, neocortical tissue was excised from patients (would normally be disposed of during surgery) and immediately placed in sterile ice-cold oxygenated 0.22 μm-filtered artificial cerebrospinal fluid (aCSF) containing 87 mM NaCl, 2.5mM KCl, 10 mM HEPES, 1.62 mM NaH2PO4, 25 mM D-glucose, 129.3 mM sucrose, 1 mM Na-Pyruvate, 1 mM ascorbic acid, 7 mM MgCl2 and 0.5 mM CaCl2. The tissue was then sub-dissected and mounted in 2% agar, before being glued (cyanoacrylate, Loctite) to a vibratome stage. 300 µm-thick slices were then cut in ice-cold and oxygenated aCSF, before being sub-dissected into smaller slices. Slices were placed into 0.22 μm-filtered wash buffer composed of oxygenated Hanks Balanced Salt Solution (HBSS, ThermoFisher: 14025092), HEPES (20 mM) and 1X penicillin-streptomycin (ThermoFisher: 15140122) (305 mOsm, pH 7.3) for 15 minutes at room temperature. Slices were then plated on membranes (Millipore: PICM0RG50) sitting on top of 750 μl of a second wash medium, placed inside 35 mm culture dishes. The second wash medium was 0.22 µm-filtered and composed of BrainPhys Neuronal Medium (StemCell Technologies: 5790) (96%), N2 (ThermoFisher: 17502001) (1X), B27 (ThermoFisher: 17504044) (1X), hBDNF (StemCell Technologies: 78005) (40 ng/ml), hGDNF (StemCell Technologies: 78058) (30 ng/ml), Wnt7a (Abcam: ab116171) (30 ng/ml), ascorbic acid (2 μM), dibutyryl cAMP (APExBIO: B9001) (1 mM), laminin (APExBIO: A1023) (1 ug/ml), penicillin/streptomycin (ThermoFisher: 15140122) (1X), nystatin (Merck: N1638) (3 units/ml) and HEPES (20 mM). Slice cultures were kept in the second wash medium in an incubator at 37°C with 5% CO2 for 1 hour, after which the medium was aspirated and replaced with maintenance medium. The maintenance medium composition was identical to that of the second wash medium, but without HEPES. 100% medium exchanges occurred twice weekly thereafter. Cultures were kept for 2 weeks and treated with either 10 μM WZ4003 (APExBIO: B1374) or DMSO (Sigma-Aldrich: D2438) control applied in culture medium, and with every feed thereafter from 0 days *in vitro (div)*. To assess neuronal integrity in HBSCs, 14 *div* slices were fixed overnight in 4% PFA, washed 3x in PBS, blocked for 1 hour in PBS +3% normal goat serum +0.5% Triton X-100 then incubated overnight at 4°C in 1/500 guinea pig anti-MAP2 (Synaptic systems: 188004) in blocking solution. The slices were then washed x3 in PBS, before incubation for 2 hours in 1/500 secondary antibody (goat anti guinea pig-488 (Thermo-Fisher: A11073)) in block buffer at room temperature. Slices were washed 3x in PBS, counterstained with DAPI for 15 minutes, washed 3x in PBS then mounted on slides in Vectashield Antifade mounting medium (2B Scientific: H-1900). Images were taken using a 63x 1.4 NA objective on a Leica TCS confocal microscope.

### Western blots

MOBSCs or HBSCs were removed from the culture membrane with a scalpel blade into RIPA buffer (ThermoFisher Scientific: 89901) with protease inhibitor cocktail (1X) and EDTA (1X) (ThermoFisher Scientific: 78429). Slices were thoroughly homogenised via trituration through 20 pipette fill and empty cycles (using a 100 μl pipette tip). 50 μl of RIPA buffer was used per MOBSC slice and 100 μl of RIPA per HBSC slice. RIPA-buffered MOBSCs, HBSC samples, or human post-mortem total protein/ synaptoneurosome samples were then mixed into equal volumes of 2X Laemmli buffer (Merck: S3401- 10VL) and boiled for 10 minutes at 98 °C. 12 µl of each sample was loaded into 4-12% NuPage Bis-Tris gels (Invitrogen: NP0336BOX), before proteins were separated by electrophoresis using MES SDS running buffer (Invitrogen: NP0002). Proteins were then transferred onto PVDF transfer membranes (Invitrogen: IB24002), before a total protein stain (Li-Cor Biosciences: 926-11016) image was acquired using a Li-Cor Odyssey Fc machine. Membranes were de-stained and then blocked for 1 hour using PBS Intercept Blocking Buffer (Li-Cor Biosciences: 927-70001). Primary antibodies were diluted in PBS Intercept Blocking Buffer with 0.1% Tween-20 and incubated with membranes overnight at room temperature, with shaking. Membranes were washed three times for 5 minutes with PBS-Tween, then incubated in darkness for 2 hours with IRDye anti-rabbit (Li-Cor Biosciences: 680RD) and anti-mouse (Li-Cor Biosciences: 800CW) secondary antibodies, each at 1:10,000 concentration. Membranes were washed 3X in PBS-Tween, 1X in PBS and then imaged using a Li-Cor Odyssey Fc machine. The following primary antibodies were used: 1:500 mouse Tau-5 (Abcam: AB80597), 1:1000 rabbit ps356 tau (Abcam: AB75603), 1:500 rabbit PSD-95 (Abcam: AB18258), 1:2500 rabbit Tuj-1 (Sigma: T2200), and 1:2000 rabbit cyclophilin-B (Abcam: AB16045). Western blot images were analysed using Empiria Studio (Version 2.3).

### Statistics

All data was analysed using R (v 4.2.2) and R Studio (v 2023.03.1, Build 446). Statistical tests were chosen according to the experimental design and dataset type. Unpaired T-tests, ratio paired T-tests and N-way repeated-measures ANOVA tests (*car* package) were conducted using linear mixed effects models (LMEM) using the *lme4* package (each model is listed in the relevant results section). Human or mouse case was included as a random effect in linear mixed effects models to avoid pseudoreplication of the data^41^. T-tests used Satterthwaite’s method, whilst Type-III F-Wald ANOVA tests used the Kenward-Roger method, to compute degrees of freedom. For N-way ANOVA analyses resulting in significant interaction effects or significant main effects where more than two within-factor levels existed, multiple comparison tests using the Tukey method for adjusting p values were computed with the *emmeans* package. Model assumptions were tested using Shapiro-Wilk and F tests, along with plotting model residuals with the ggplot2 package. To assess equality of variance, standardised model residual scatter was plotted against fitted values. Residual distribution was assessed for normality by plotting a histogram and kernel density estimate of standardised model residuals over a standard normal distribution, and by generating a quantile-quantile plot of dataset quantiles against normal quantiles. Statistical tests performed on datasets represented in Figures 5-7 were conducted on absolute data that was log-transformed, in order to compute ratio differences between differentially treated samples from the same animal/human and to satisfy assumptions of normality. Figures 5-7 represent control-normalised data to display within-animal and within-human case effects. Significance values are reported as *p* < 0.05 *, *p* < 0.01 *, *p* < 0.001 *** and error bars represent the mean ± SEM.

## Results

### p-tau Ser356 increases in a Braak Stage-dependent manner in human post-mortem temporal cortex

To investigate how the levels of p-tau Ser356 change with Braak stage in post-mortem human temporal cortex (BA20/21), total homogenate and synaptoneurosome preparations were generated from frozen post-mortem brain from 10 Braak 0-I individuals, 10 Braak III-IV individuals and 10 Braak VI individuals (with clinically diagnosed AD). Individual patient details are listed in **Table 1** with summary demographic information listed in **Table 2**. Levels of p-tau Ser356 and total tau (tau5) were quantified by Western blot **(Fig. 2a).** Total tau (tau5) was detected in all cases, and was normalised to total protein (REVERT protein stain). For statistical analysis, the following LMEM was applied: *Protein level ∼ Braak Stage * Preparation + (1|CaseID).* The levels of total tau did not increase with Braak Stage (**Fig. 2b**, effects of Braak Stage: F(2,46.01)=1.32, p=0.28) and there were no detectable differences in the levels of tau in total homogenate compared to synaptoneurosome preparations (**Fig. 2b**, effect of preparation: F(1,27.00)=0.15, p=0.71). By contrast, the levels of p-tau Ser356, normalised to total tau levels, showed a significant increase with Braak stage (**Fig. 2c** effect of Braak stage: **F(2,42.28)=5.72, p=0.006) and a detectable effect of preparation (**Fig. 2c**, effect of preparation: *F(1,27.00)=4.94, p=0.035) driven by an increase in p-tau Ser356 in total homogenate, compared to synaptoneurosome, in Braak VI cases (**Fig. 2c**, multiple comparisons test: *p=0.034), possibly reflective of increased tau in the cell bodies at a stage where tangle prevalence will be high. There was a significant increase in p-tau Ser356 in Braak VI AD brains compared to Braak 0-I control brains (**Fig. 2c**, multiple comparisons test:***p=0.0009), and a strong trend for increase between Braak III-IV and Braak VI (**Fig. 2c**, multiple comparisons test: p=0.053).

**Figure 2:**
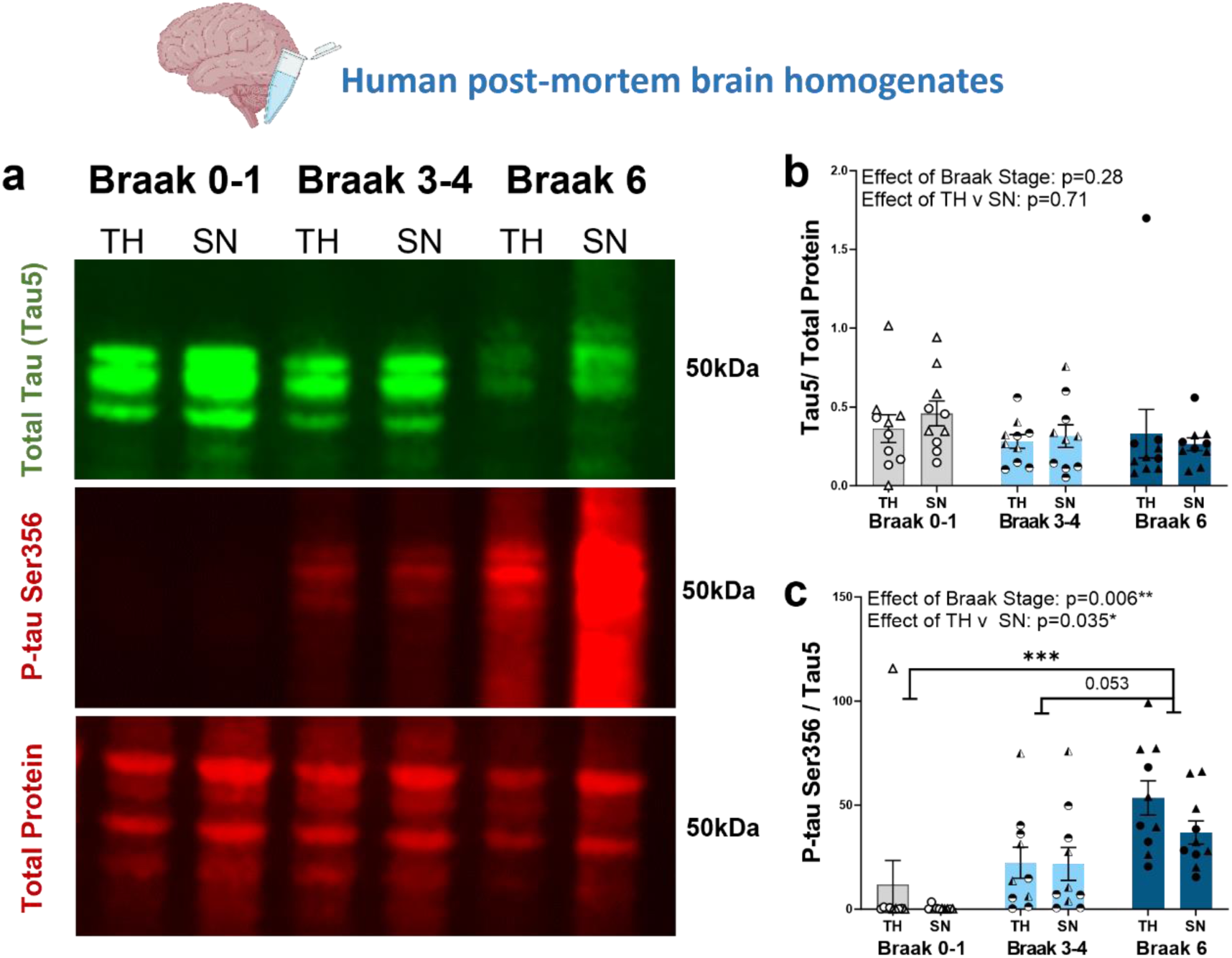
ps356 Tau Expression Increases in a Braak Stage-Dependent Manner in post-mortem AD Temporal Cortex. ***(a)** Representative Western blot image of total homogenates (TH) and synaptoneurosomes (SN) from Braak 0-1, 3-4 or 6 (diagnosed AD) stage post-mortem brain. Blots were probed for total protein (REVERT stain), total tau (Tau5) and p-Tau Ser356. **(b)** There is no significant impact of Braak Stage (F_(2,46.01)_=1.32, p=0.28) or preparation (TH v SN) (F_(1,27.00)_=0.15, p=0.71) on the levels of total tau (Tau 5) in human postmortem brain. **(c)** There is a significant increase in P-tau Ser356 (normalised to total tau) with increasing Braak Stage (**F_(2,42.28)_=5.72, p=0.006), with a significant difference between Braak 0-I and Braak VI AD (***T_(27)_=4.13, p=0.0009) and a strong trend increase between Braak III-IV and Braak VI (T_(27)_=2.45, p=0.053). There is an effect of preparation (*F_(1,27.00)_=4.94, p=0.035), with increased p-tau Ser356 in the total homogenate relative to synaptoneurosomes in Braak VI AD post-mortem brains (*T_(27)_=2.23, p=0.034). Each point on the graphs represents a single case, triangles= males, circles= females. n = 10 cases per Braak Stage. Cartoons generated using BioRender.*

### Co-localisation of p-tau Ser356 with ThioS positive tangles and dystrophic neurites in AD post-mortem brain tissue

Having established a significant increase in p-tau Ser356 in AD brains, we sought to establish where in the brain tissue this form of tau is located and its inclusion in neurofibrillary tangles (NFTs). Paraffin sections were obtained from 5 control (Braak 0-II) and 5 confirmed AD (Braak VI, Thal 5) cases (individual patient details listed in **Table 1**, summary demographic details listed in **Table 3**). Staining in paraffin sections demonstrated that p-tau Ser356 is readily detectable in tangles, dystrophic neurites, and some reactive astrocytes in AD brain (**Fig. 3**). Of the 85 ThioS positive tangles we identified in the AD cases, 79 were positive for p-tau Ser356 (93%) (**Fig. 3f**). Interestingly, whilst control brains showed, as expected, significantly lower levels of neuropathology (**Fig. 3a**) we also found evidence of p-tau Ser356 staining in some control brains, in areas co-localising with ThioS, most notably in dystrophic neurites (**Fig. 3b**). Together, these results suggest that p-tau Ser356 is a common component of dystrophic neurites and Thio-S positive NFTs, and may therefore be involved early in the tau aggregation cascade.

**Figure 3:**
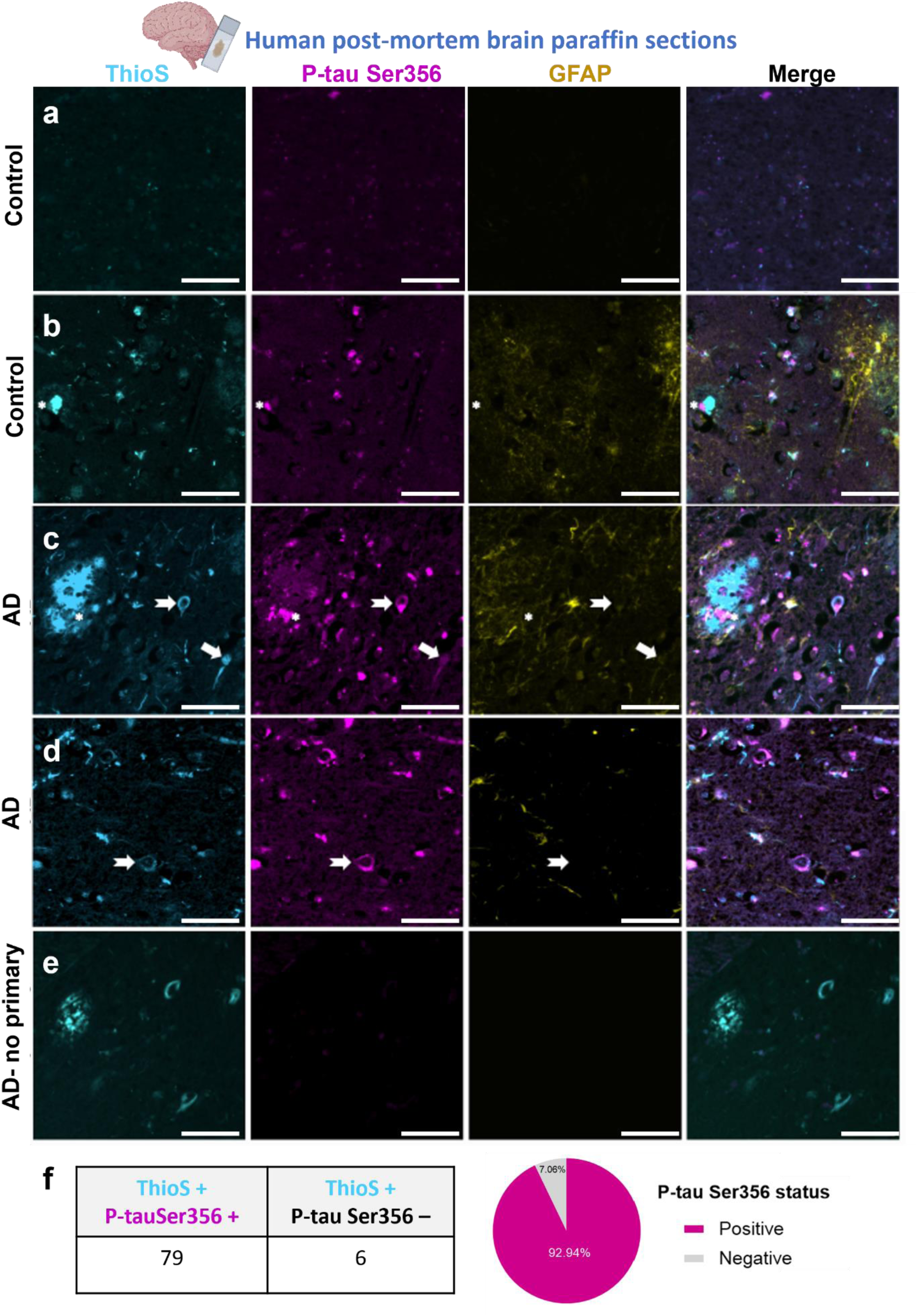
P-tau Ser356 is observed around plaques, in dystrophic neurites and in tangles in human post-mortem brain. *Paraffin sections of control (**a,b)** or AD **(c-e)** post-mortem brain stained for plaques/tangles (ThioS-cyan), p-tau Ser356 (magenta), and reactive astrocytes (GFAP-yellow). In human brain sections, pTau356 (magenta) is found in tangles (arrows) and dystrophic neurites around plaques (asterisks). In tangles, p-tau Ser356 is both seen labelling Thioflavin S positive fibrils (arrows) and in punctate patterns in neurons containing Thioflavin S positive tau fibrils (notched arrows). pTau356 is also observed in some GFAP positive astrocytes (yellow). Control brain with little/no ThioS or p-tau Ser356 staining **(a).** Control brain with evidence of dystrophic neurites **(b)**, AD brains with evidence of plaques **(c)**, dystrophic neurites **(c)** and tangles **(c, d)**. No primary antibody control **(e)**. Scale bar 50μm. **(f)** Quantification of ThioS, p-tau Ser356 double positive tangles, compared to ThioS only tangles shows the vast majority of tangles counted in this study (93%) contain p-tau Ser356. Cartoons generated using BioRender.*

**Table 3:** Summary demographic information of human post-mortem subjects (Paraffin sections study (Figure 3)). Cartoons generated using BioRender.

|  | Control<br>(N=5) | AD<br>(N=5) | Overall<br>(N=10) |
| --- | --- | --- | --- |
| **Age** |  |  |  |
| Mean (SD) | 79.4 (2.70) | 80.6 (9.07) | 80.0 (6.34) |
| Median [Min, Max] | 79.0 [77.0, 84.0] | 85.0 [66.0, 89.0] | 79.0 [66.0, 89.0] |
| **Sex** |  |  |  |
| F | 3 (60.0%) | 2 (40.0%) | 5 (50.0%) |
| M | 2 (40.0%) | 3 (60.0%) | 5 (50.0%) |

### p-tau Ser356 co-localises with pre-synaptic terminals in AD post-mortem brain tissue

Given recent work highlighting the role of synaptic tau in the early stages of AD pathology^22^ we next sought to examine the presence of p-tau Ser356 at synaptic terminals. Using array tomography, which physically overcomes the diffraction limitations of light in the axial plane to enable imaging of the protein composition of individual synapses^29^, we imaged 5 confirmed AD (Braak V-VI) and 5 age/sex matched control cases (Braak 0-III) for p-tau Ser356, AT8 (detects tau phosphorylated at Ser202 & Thr205) and the pre-synaptic marker synaptophysin (**Fig. 4a,b**). Individual patient demographic data is listed in **Table 1** and summary demographic data listed in **Table 4**. For statistical analysis, the following LMEM was applied: *Co-localisation % ∼ Diagnosis + Sex + (1|CaseID)*. Whilst largely undetectable in control cases (**Fig. 4a**), both p-tau Ser356 (**Fig. 4c**) and AT8 (**Fig. 4d**) were found to co-localise with synaptophysin in AD cases (**t(7.14)=4.32, p=0.0033 and **t(7.13)=3.82, p=0.0063 respectively). Interestingly, whilst p-tau Ser356 and AT8 were found to co-localise with a similar percentage of pre-synaptic terminals (median of 1.38% (**Fig. 4c**) and 1.60% (**Fig. 4d**) respectively), the percentage of synaptophysin puncta that co-localised with both tau epitopes, (whilst still significantly higher in AD than controls (*t(7.08)=3.242, p=0.014)), was considerably smaller (median of 0.36% (**Fig. 4e**)). This could suggest that different synapses have unique signatures of tau phosphorylation patterns in AD, or that there are technical considerations (e.g. masked antibody binding sites) that make it harder to resolve both epitopes when they co-localise. 3D reconstruction images (**Fig. 4f-i**) show examples of synaptic co-localisation either AT8 alone (**Fig. 4g**), p-tau Ser356 alone (**Fig. 4h**) or both tau epitopes together (**Fig. 4i**).

**Figure 4:**
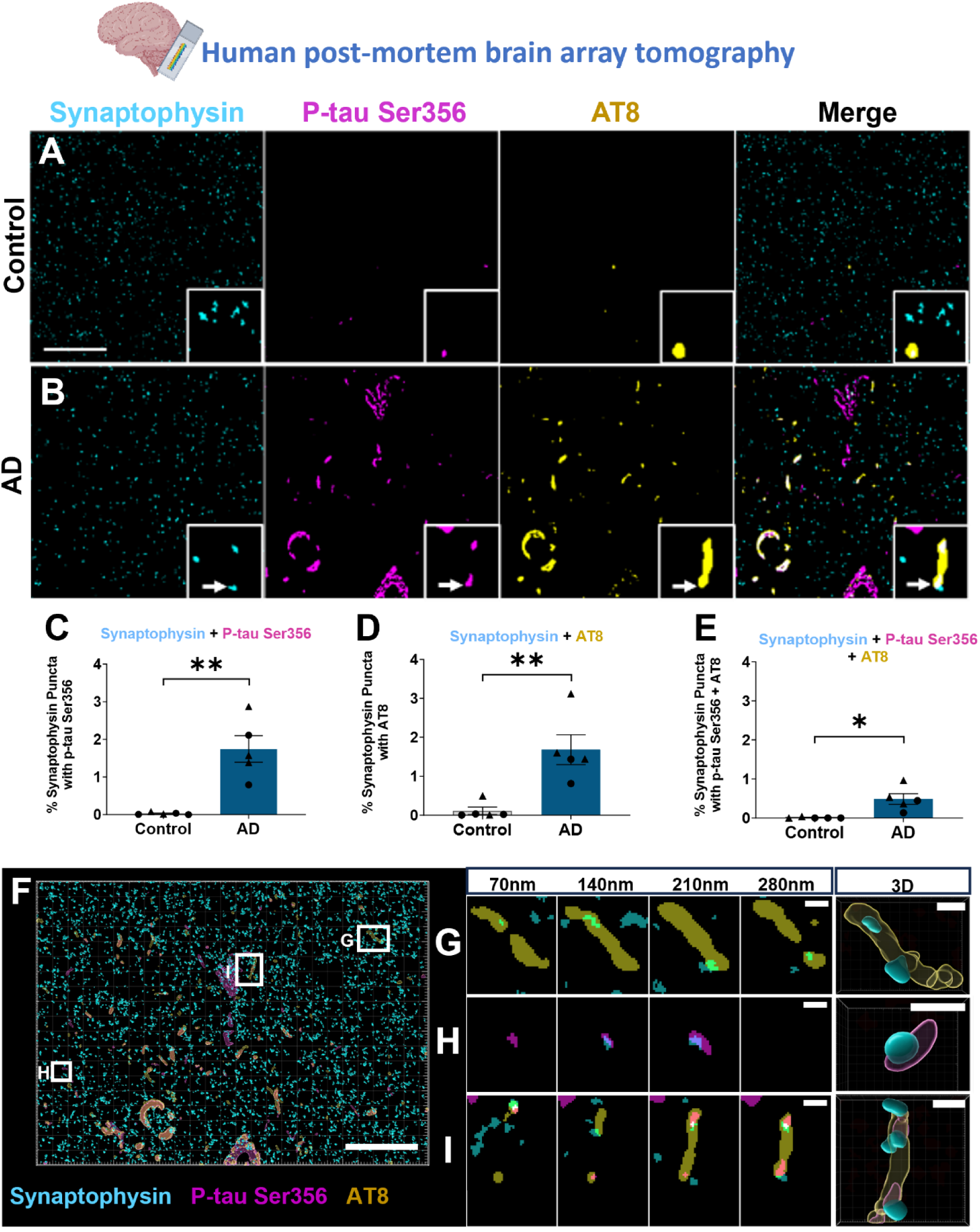
p-tau Ser356 co-localises at the synapse in Alzheimer’s disease post-mortem brain. *(**a**,**b**) Representative images of 70nm thick array tomography images from control **(a)** or AD **(b)** post-mortem brain. Sections were stained with the presynaptic marker synaptophysin (cyan), p-tau Ser356 (magenta), AT8 (yellow). The white arrow indicates the area of co-localisation of synaptophysin, p-tau Ser356 and AT8 in the AD brain case. Scale bar represents 20µm. There is a significant increase in synaptophysin co-localising with p-tau Ser356 in AD brain (**T_(7.14)_=4.324, p=0.0033) **(c).** There is a significant increase in synaptophysin colocalising with AT8 in AD brain (**T_(7.13)_=3.825, p=0.0063) **(d)** and a significant increase in synapses containing both p-tau Ser356 and AT8 (*T_(7.08)_=3.242, p=0.014) **(e).** Each point on the graph represents a single case, males= triangles, females= circles. n = 5 control and 5 AD cases. Representative image and 3D constructions from serial 70nm section from AD brain **(f-i)** showing synaptophysin (cyan) co-localisation with AT8 (yellow) **(g)**, p-tau Ser356 (magenta) **(h)** and synapses co-localising with both AT8 and p-tauSer356 **(i)**. Scale bar represents 25µm in **f** and 1µm in **g-i**. Cartoons generated using BioRender.*

**Table 4:**
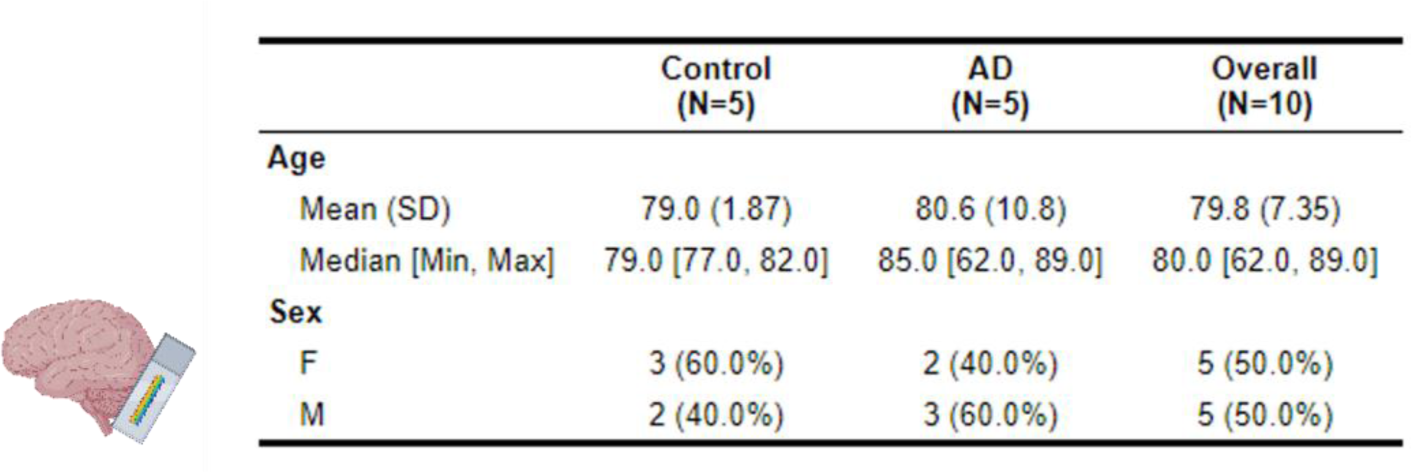
Summary demographic information of human post-mortem subjects (Array tomography study (Figure 4)). Cartoons generated using BioRender.

### The NUAK inhibitor WZ4003 lowers p-tau Ser356 and total tau in MOBSCs in a culture phase dependent manner

Having confirmed that p-tau Ser356 correlates with AD Braak stage, is located in the majority of NFTs, and is also found in a subset of presynaptic terminals, we sought to evaluate potential pharmacological tools to lower p-tau Ser356 in live brain tissue. To determine whether response to NUAK inhibition differs under physiological conditions versus conditions of early amyloid dysregulation (which may initiate downstream tau changes) we generated mouse organotypic brain slice cultures (MOBSCs) from wildtype and APP/PS1 littermates. We sought to assess whether NUAK inhibition has different outcomes depending on the timing of treatment application. In previous work in MOBSCs, it has been noted that there is inflammation, reorganisation and recovery in the first 2 weeks *in vitro*, with cultures stabilising after this time^33,42,43^. MOBSCs were generated from WT or APP/PS1 littermate P6-P9 mice such that four separate culture dishes were generated per animal (**Fig. 5a**). Cultures were split into two culture phase groups (0-2 weeks *in vitro*, vs 2-4 weeks *in vitro*) and two treatment groups (DMSO or WZ4003) resulting in four experimental conditions represented in tissue from the **same animal**: 0-2 weeks DMSO, 0-2 weeks WZ4003, 2-4 weeks DMSO, 2-4 weeks WZ4003. 0-2 week cultures were treated at 0 *div*, then harvested for protein analysis at 14 *div*. 2-4 week cultures were left untreated for the first 2 weeks in culture, before undergoing treatment at 2 weeks *in vitro* and being harvested at 4 weeks *in vitro* for processing via Western blot (**Fig. 5b**). In accordance with prior literature^9^, we applied 5 µM WZ4003 to cultures. For graphical purposes, all conditions are displayed as normalised to the 0- 2 week DMSO-treated culture from the same animal, whilst analysis was performed using ratio repeated measured analysis with the absolute data (Graphs of absolute data are displayed in **Supp. Fig. 1**). For statistical analysis, the following LMEM was applied: *Protein level ∼ Genotype * Phase * Treatment + (1|Litter/Animal)*.

**Figure 5:**
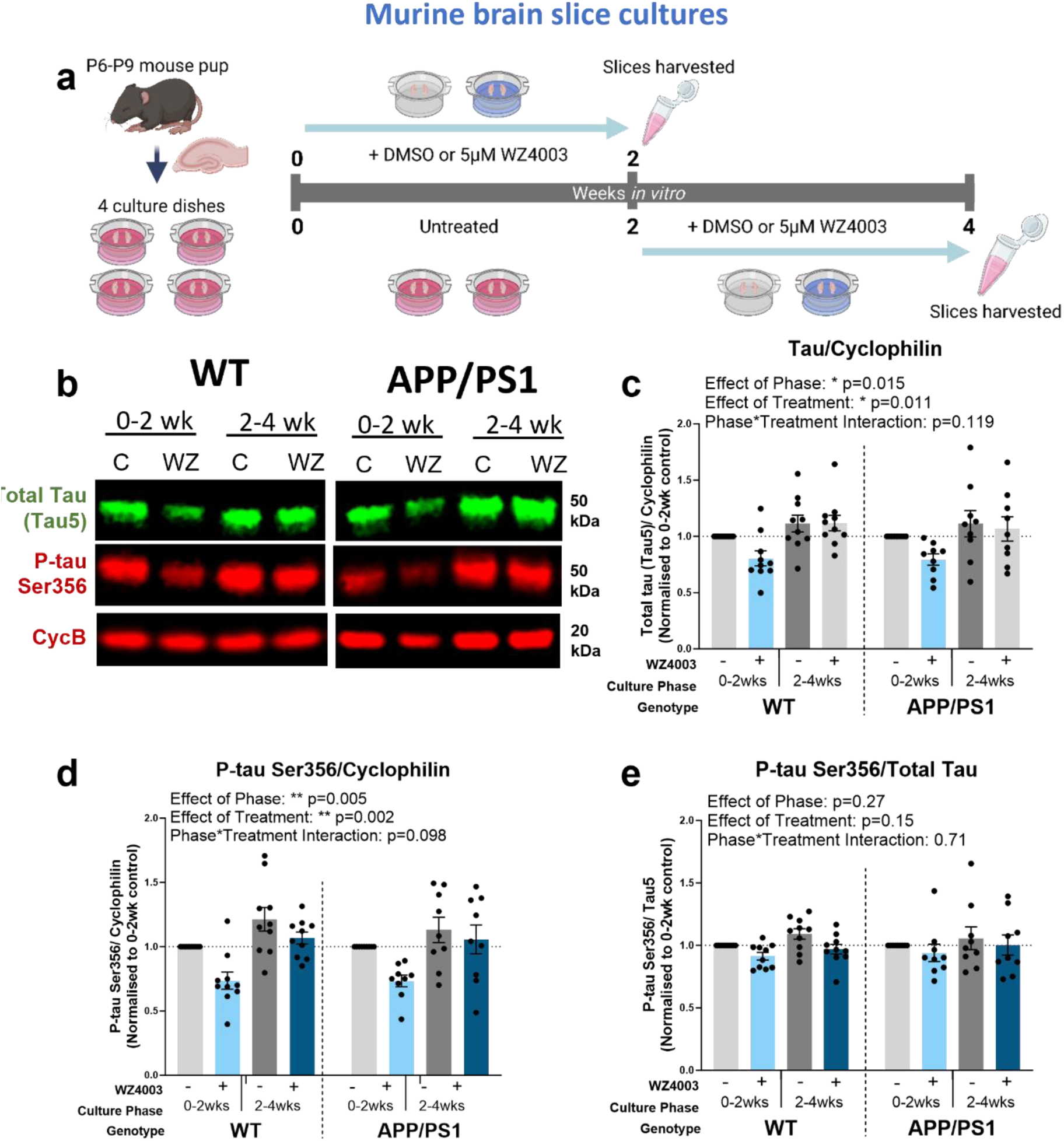
WZ4003 lowers tau and phospho-tau in a phase dependent manner in MOBSCs. (a) *Schematic showing the dissection and treatment schedule for MOBSCs. 4 dishes are generated per animal, split into 4 treatment conditions: 0-2 weeks control, 0-2 weeks WZ4003, 2-4 weeks control, 2-4 weeks WZ4003. Slices are harvested for Western blot at the end of their treatment phase and ratio change in protein levels (normalised to cyclophilin B) compared within samples from the same mouse. **(b)** Representative Western blot for total tau, p-tau Ser 356 and housekeeping protein cyclophilin. The following graphs are displayed normalised to 0-2 week control for each animal to show relative differences, but statistics are performed on absolute data (absolute data displayed in Supp Fig 1). **(c)** There is a significant effect of phase (*F_(1,51.00)_=6.28, p=0.015) and treatment (*F_(1,51.00)_=7.01, p=0.011) on the levels of total tau, but no effect of genotype (F_(1,21.30)_=0.02, p=0.894). **(d)** There is a significant effect of phase (**F_(1,51.00)_=8.73, p=0.005) and treatment (**F_(1,51.00)_=10.67, p=0.002) on the levels of p-tau Ser356, and a trend interaction between Phase*Treatment (F_(1,51.00)_=2.83, p=0.098), but there is no effect of genotype (F_(1,26.41)_=0.00, p=0.997). **(e)** There are no significant effects of phase (F_(1,51.00)_=1.25, p=0.27), treatment (F_(1,51.00)_=2.10, p=0.15) or genotype (F_(1,26.51)_=0.08, p=0.781) on the ps356/total tau ratio. N=9 APP/PS1 and 10 WT animals, 1-2 slices per animal per condition. Cartoons generated using BioRender.*

Regardless of genotype, we found a significant effect of culture phase on the levels of both total tau (**Fig. 5c**, effect of phase: * F(1,51.00)=6.28, p=0.015) and p-tau Ser356 (**Fig. 5d**, effect of phase: ** F(1,51.00)=8.73, p=0.005), indicating that the levels of tau protein rise over time in culture in MOBSCs, possibly reflecting the regrowth or maturation of neurites following the initial slicing procedure. The ratio of p-tauSer356/total tau remained stable over culture phase (**Fig. 5e**, effect of phase: F(1,51.00)=1.25, p=0.27). Application of WZ4003 resulted in a significant reduction in both total tau (**Fig. 5c**, effect of treatment: * F(1,51.00)=7.01, p=0.011) and p-tau Ser356 (**Fig. 5d**, effect of treatment: ** F(1,51.00)=10.67, p=0.002) with the ratio of p-tau Ser356/ total tau remaining relatively stable, indicating a largely proportional loss of both forms of tau in response to WZ4003 (**Fig. 5e**, effect of treatment: F(1,51.00)=2.10, p=0.15). Whilst there was not a significant treatment*phase interaction for total tau (**Fig. 5c**, effect of treatment*phase: F(1,51.00)=2.52, p=0.119), there was a trend interaction for p-tau Ser356 (**Fig. 5d**, effect of treatment*phase: F(1,51.00)=2.83, p=0.098), where the mean percentage loss of protein levels between treated and control samples trended to be larger in the 0-2 week phase than the 2-4 week phase. The reduction in p-tau Ser356 averaged at 26.41% for WT and 26.64% for APP/PS1 0-2 week cultures, compared with 9.13% for WT and 6.84% for APP/PS1 2-4 week cultures.

In this study, we found no differences between genotypes in either baseline levels of tau, or in the degree of response to treatment. There was no effect of genotype on the p-tau Ser356/total tau ratio (**Fig. 5e/ Supp. Fig. 1c**: effect of genotype F(1,26.51)=0.08, p=0.781), indicating that, in MOBSCs from APP/PS1 mice up to 4 weeks *in vitro,* the presence of amyloid mutations does not result in perturbations to p-tau Ser356.

### Synaptic and neuronal proteins rise over time in culture in MOBSCs, and are impacted by WZ4003 treatment

We next sought to examine the impact of WZ4003 treatment on both synaptic and neuronal protein levels in MOBSCs (**Fig. 6**). For statistical analysis, the following LMEM was applied: *Protein level ∼ Genotype * Phase * Treatment + (1|Litter/Animal).* Westen blot analysis (**Fig. 6a**) revealed that the synaptic protein PSD95 was upregulated in 2-4 week cultures compared to 0-2 week cultures (**Fig. 6b**, effect of phase: *** F(1,51.00)=26.65, p<0.001), but, interestingly, was *reduced* in response to WZ4003 (**Fig. 6b**, effect of treatment: * F(1,51.00)=5.60, p=0.022). The neuronal tubulin marker Tuj1 is similarly upregulated in 2-4 week cultures (**Fig. 6c**, effect of phase: *** F(1,51.00)=13.36, p<0.001), but was not significantly impacted by WZ4003 treatment (**Fig. 6c**, effect of treatment: F(1,51.00)=1.99, p=0.164). To examine the impact of synaptic proteins in the context of changes to neuronal proteins, we next normalised the levels of PSD95 to Tuj1 levels. Interestingly, the effect of phase remains (**Fig. 6d**, effect of phase: * F(1,51.00)=4.39, p=0.041), indicating that synaptic proteins rise in excess of the rise in neuronal proteins over time, but the effect of treatment disappears (**Fig. 6d**, effect of treatment: F(1,51.00)=1.63, p=0.207). This suggests that, despite the loss of Tuj1 in response to WZ4003 treatment not reaching significance alone, a WZ4003-induced loss of neuronal protein may partly contribute to the loss of PSD95. There were no differences between genotypes for either PSD95 or Tuj1 at baseline or in response to treatment (**Fig. 6, Supp. Fig. 1**).

**Figure 6:**
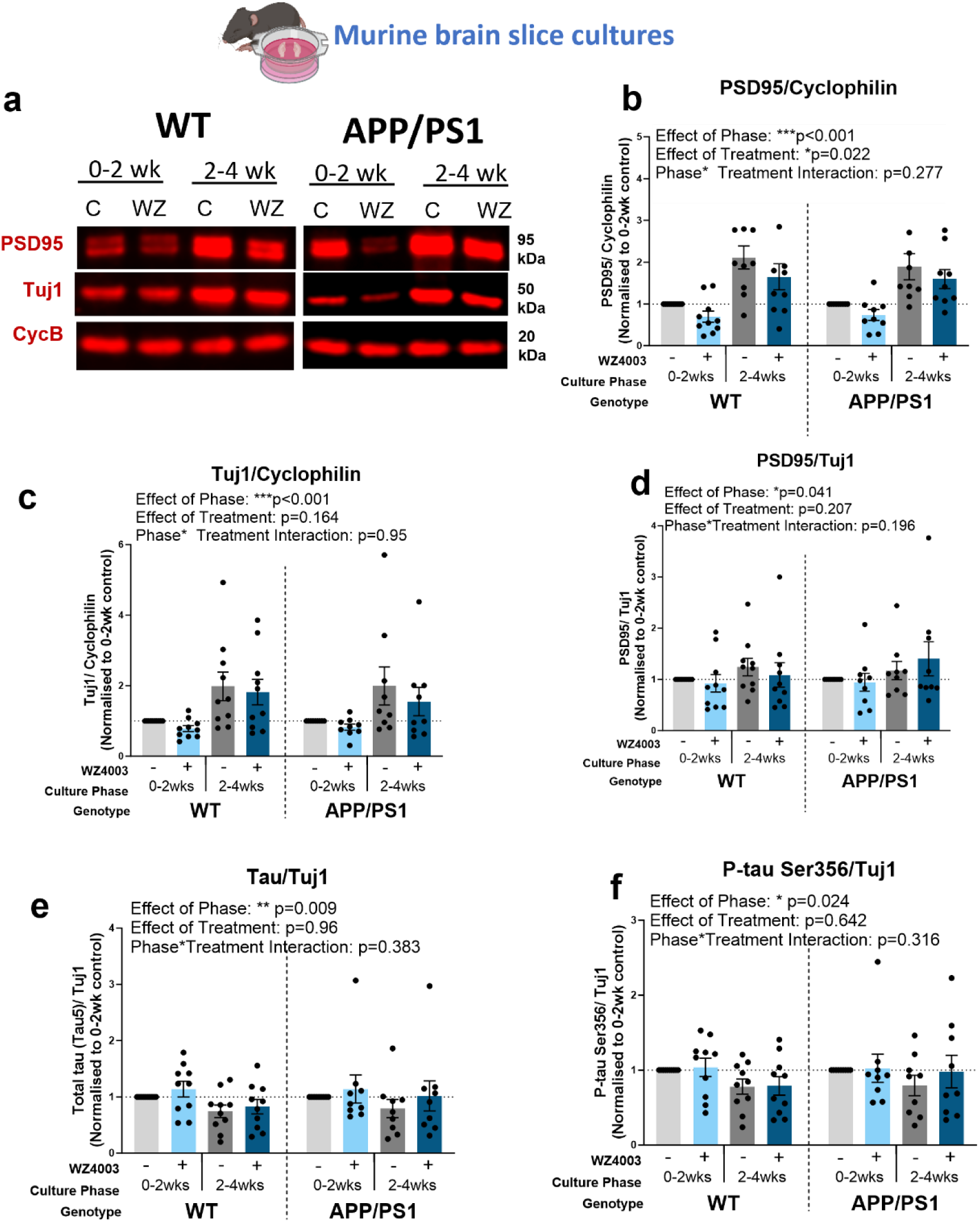
Neuronal and synaptic proteins increase over time in culture and are impacted by WZ4003 treatment. ***(a)** Representative Western blot for PSD95, Tuj1 and housekeeping protein cyclophilin. The following graphs are displayed normalised to 0-2 week control for each animal to show relative differences, but statistics are performed on absolute data (absolute data displayed in Supp Fig 1). **(b)** There is a significant effect of phase (***F_(1,51.00)_=26.65, p<0.001) and treatment (*F_(1,51.00)_=5.60, p=0.022) on the levels of PSD95 normalised to cyclophilin, but no effect of genotype (F_(1,41.92)_=0.21, p=0.652). **(c)** There is a significant effect of phase (***F_(1,51.00)_=13.36, p<0.001) on the levels of Tuj1 normalised to cyclophilin, but no effects of treatment (F_(1,51.00)_=1.99, p=0.164) or genotype (F_(1,41.92)_=0.00, p=0.957). **(d)** There is a significant effect of phase (*F_(1,51.00)_=4.39, p=0.041) but no effect of treatment (F_(1,51.00)_=1.63, p=0.207) or genotype (F_(1,47.89)_=0.13, p=0.715) on the levels of PSD95 when normalised to Tuj1. **(e)** There is a significant effect of phase (**F_(1,51.00)_=7.38, p=0.009), but no effect of treatment (F_(1,51.00)_=0.00, p=0.964) or genotype (F_(1,42.38)_=0.08, p=0.774), in the levels of total tau when normalised to Tuj1. **(f)** There is a significant effect of phase (*F_(1,51.00)_=5.40, p=0.024), but no effect of treatment (F_(1,51.00)_=0.22, p=0.642) or genotype (F_(1,42.83)_=0.00, p=0.957), on the levels of p-tau Ser356 when normalised to Tuj1. N=9 APP/PS1 and 10 WT animals, 1-2 slices per animal per condition. Cartoons generated using BioRender.*

When normalising total tau (**Fig. 6e**) or p-tau Ser356 (**Fig. 6f**), to Tuj1, we see that, relative to neuronal tubulin, there is a *reduction* in the levels of total tau (**Fig, 6e**, effect of phase: ** F(1,51.00)=7.38, p=0.009) and p-tau Ser356 (**Fig. 6f**, effect of phase= * F(1,51.00)=5.40, p=0.024) in the 2-4 week cultures. This indicates, whilst there may be increased neuronal (**Fig. 6c**) and tau (**Fig 5c,d**) protein over time, the *proportion* of tau relative to neuronal protein declines as the cultures age. Of particular note, is that the effect of WZ4003 treatment is abolished when normalising total tau (**Fig. 6e**, effect of treatment: F(1,51.00)=0.00, p=0.964), or p-tau Ser356 (**Fig. 6f**, effect of treatment: F(1,51.00)=0.22, p=0.642) to Tuj1, demonstrating that the lowering of tau, inhibition of NUAK, or other impacts of WZ4003 results in a proportional reduction in both tau and neuronal protein in MOBSCs up to 4 weeks *in vitro*.

### WZ4003 alters p-tau Ser356 levels in live human brain slice cultures

Finally, we sought to assess the impact of WZ4003 treatment in live adult human brain tissue. Healthy, peri-tumoral access tissue from 5 patients (demographics details listed in **Table 5**), was processed into 300 μm slices and cultured on membrane inserts for 2 weeks *in vitro* (**Fig. 7a**). Cultures showed intact MAP2 positive neuronal cell bodies and processes at 14 *div* (**Fig. 7b**) and tau, neuronal and synaptic proteins were readily detectable by Western blot (**Fig. 7c**). Slices from the same individuals were divided into both control (DMSO) and treatment (10 μM WZ4003) conditions, and protein levels were compared to the untreated control from the same patient. Graphs showing control-normalised data are used for display purposes (**Fig. 7**), with statistics run on ratio paired t-tests using absolute data. Graphs of absolute data are shown in **Supp. Fig. 2**. For statistical analysis, the following LMEM was applied: *Protein level ∼ Treatment + (1|CaseID)*.

**Figure 7:**
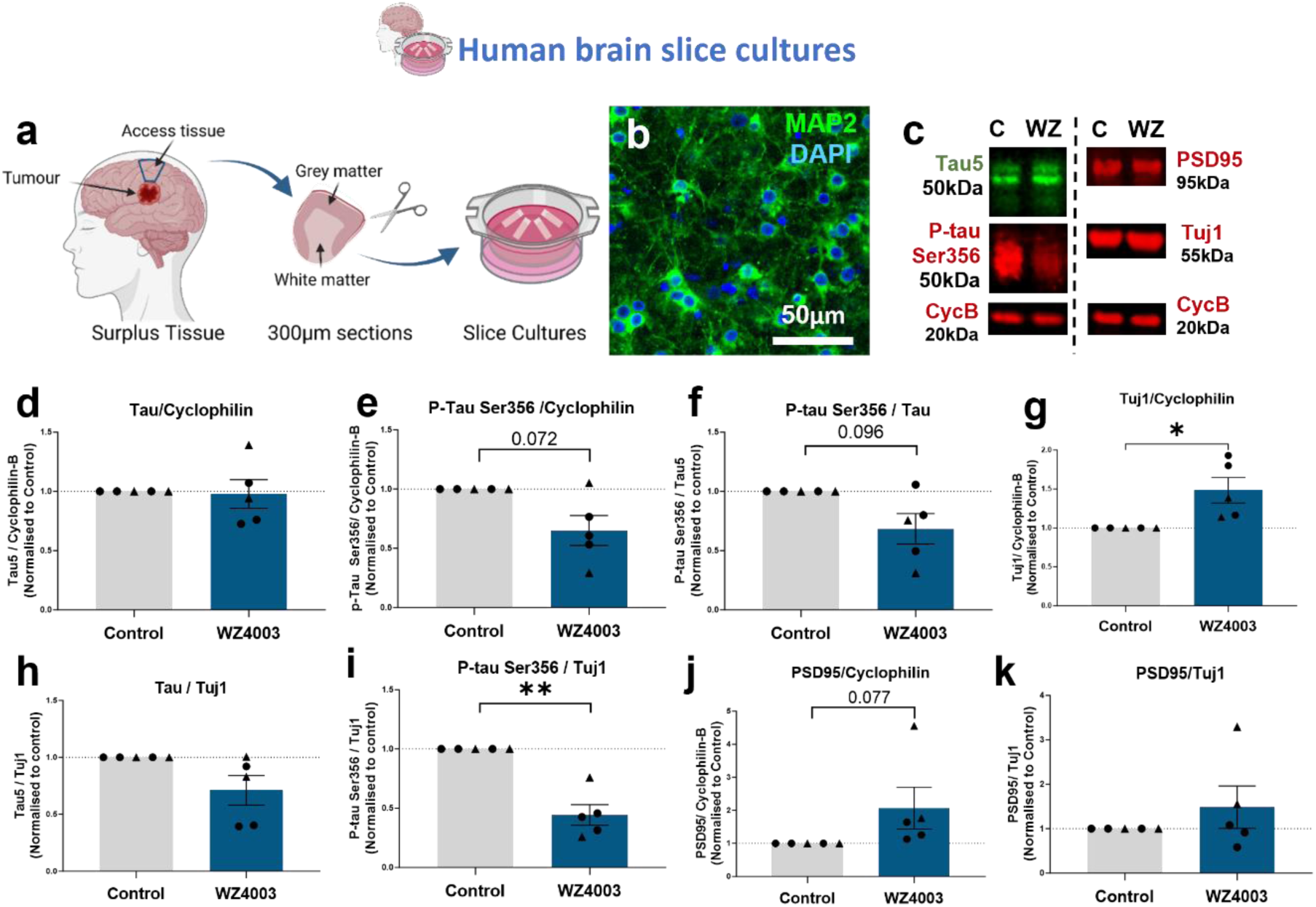
Human slice cultures are responsive to WZ4003 treatment. ***(a)** Cartoon illustrating the work-flow for generating human brain slice cultures (HBSCs) from surplus access tissue from neurosurgical procedures. **(b)** MAP2 (green) and DAPI (blue) staining shows intact neuronal cell bodies and neurite processes in 14div HBSC (scale bar= 50μm). **(c)** Representative Western blot from 14div HBSC showing total tau (tau5), p-tau Ser356, PSD95, Tuj1 and housekeeping protein cyclophilin B. **(d)** WZ4003 does not significantly alter levels of tau (normalised to cyclophilin) (T_(4)_=0.43, p=0.688). **(e)** There is a trend for WZ4003 treatment to reduce p-tau Ser356 (normalised to cyclophilin) (T_(4)_=2.43, p=0.072). **(f)** There is a trend for WZ4003 to reduce the ratio of p-tau Ser356/ total tau (T_(4)_=2.17, p=0.096). **(g)** WZ4003 treatment significantly increased Tuj1 levels (*T_(4)_=3.39, p=0.028). **(h)** WZ4003 does not significantly alter tau levels as a proportion of neuronal protein (T_(4)_=2.04, p=0.11). **(i)** WZ4003 significantly lowers p-tau Ser356 as a proportion of neuronal protein (**T_(4)_=4.81, p=0.0086). **(j)** There is a trend for WZ4003 to increase PSD95 protein (normalised to cyclophilin) (T_(4)_=2.37, p=0.077). **(k)** There is no effect of WZ4003 on PSD95 protein (normalised to Tuj1) (T_(4)_=0.74, p=0.501). N=5 cases per condition. Each point represents an individual human case, triangles= males, circles= females. Cartoons generated using BioRender.*

**Table 5:**
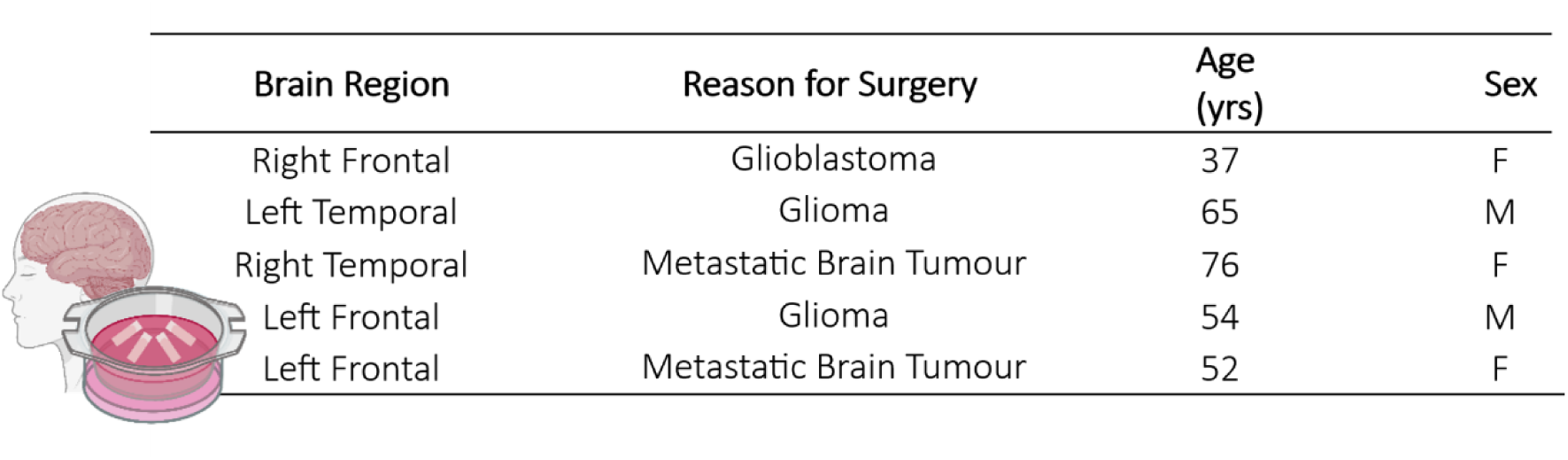
Demographic, brain region and reason for surgery information for human samples obtained from neurosurgical procedures. Tissue obtained from these individuals went on to generate the HBSCs used in Figure 7. Cartoons generated using BioRender.

We found that WZ4003 treatment did not impact the level of total tau when normalised to the housekeeping protein cyclophilin (**Fig**. **7d**, mean % decrease= 2.21%, T(4)=0.43, p=0.688), but there was a trend for reduced levels of p-tau Ser356 (**Fig. 7e**, mean % decrease = 35.03%, T(4)=2.43, p=0.072) and a trend for reduced p-tau Ser356/tau ratio (**Fig. 7f**, mean % decrease = 31.63%, T(4)=2.17, p=0.096). Interestingly, in contrast to the response in MOBSCs, we found a significant *increase* in the levels of the neuronal protein Tuj1 (**Fig. 7g**, mean % increase = 89.15%, * T(4)= 3.39, p=0.028). When normalising tau levels to Tuj1, we saw no significant loss of total tau (**Fig. 7h**, mean % decrease = 29%, T(4)= 2.04, p=0.11), but a significant loss of p-tau Ser356 (**Fig. 7i**, mean % decrease = 55.73%, ** T(4) = 4.81, p=0.0086), indicating that WZ4003 treatment results in preferential lowering of p-tau Ser356 without incurring a loss of neuronal protein in HBSCs. We saw a trend for increased PSD95 in the WZ4003 treated cultures (**Fig. 7j**, mean % increase = 104.88%, T(4)=2.37, p=0.077), that was likely proportional to the rise in Tuj1 levels (**Fig. 7k**, T(4)=0.739, p=0.501). Overall, this demonstrates that WZ4003 treatment specifically reduces p-tau Ser356 in adult human brain tissue, relative to increases in neuronal and synaptic proteins.

## Discussion

This work combines multiple experimental tools (human post-mortem, mouse organotypic brain slice, human organotypic brain slice (**Fig. 1c**)) to better characterise the timing and location of p-tau Ser356 in AD and assess the impact of pharmacological NUAK inhibition under a number of physiological and pathological conditions. Our work highlights p-tau Ser356 as a highly disease-associated form of tau in post-mortem AD human brain. We show that p-tau Ser356 is not readily detectable in protein extracts from control post-mortem brains, but can localise to dystrophic neurites surrounding areas of sporadic pathology in control tissue paraffin sections. We find an effect of Braak stage on the accumulation of p-tau Ser356 in post-mortem temporal lobe (BA20/21), with increased protein levels being detectable in post-mortem brain from Braak stage III-IV cases, and the highest levels in Braak VI cases. When examining paraffin sections from AD brain, we find that almost all (93%) of ThioS positive tangles are dual-labelled with p-tau Ser356, indicating this epitope may be phosphorylated early on in the tangle formation process. This finding is in agreement with studies in Drosophila^14^ and neuroblastoma cells^9^ suggesting that phosphorylation at this site can promote downstream phosphorylation of multiple other sites. The consistent and early appearance of p-tau Ser356 in the AD disease course, once again highlights this epitope as a potential therapeutic target.

For the first time, we used array tomography, a microscopy method permitting sub-diffraction limit resolution characterisation of protein composition of individual synapses^29^, to assess whether p-tau Ser356 is present at the synapse in AD brains. We found that, whilst p-tau Ser356 is almost undetectable in control brain synapses, there is a small but significant proportion (∼1-3%) of synapses that co-localise with p-tau Ser356 in AD brain. Interestingly, whilst a similar proportion of synapses co-localise with AT8 (p-tau Ser202/Thr205), the proportion of synapses that contain both epitopes is considerably lower (∼0-1%), raising the possibility that the order of tau phosphorylation may be different in individual synapses. Alternatively, it may be that detection of both epitopes together is under-represented through technical limitations, such as reduced antibody binding when both epitopes co-localise. Recent work has highlighted potentially important roles of synaptic tau for both toxicity and involvement of trans-synaptic tau spread^22–25^. Future work exploring the impact of synaptic p-tau Ser356, in contrast to tau phosphorylated at alternative sites, could further elucidate its role in AD pathology.

Given the potential importance of NUAK1 in the phosphorylation of tau at Ser356, and our findings that p-tau Ser356 is highly associated with disease progression in AD, we sought to characterise the impacts of pharmacological NUAK inhibition under a number of physiological and disease-model conditions. In this study, we used WZ4003, which has previously been shown to be a potent inhibitor of NUAK1, and to a lesser extent NUAK2, and with no inhibitory activity on a panel of 139 other related kinases^30^. Previous studies using WZ4003 have used simple *in vitro* systems such as primary culture^44^ or cell lines^9,30^ which may oversimplify the impacts of NUAK inhibition on brain tissue containing multiple cell types, and functionally relevant neuronal architecture^34^. In addition, prior work looking at the effect of NUAK1 knockdown in animal models (Drosophila and mouse), focused on models with tau pathology exclusively, leaving a gap in our understanding of how amyloid dysregulation, or elevated Aβ production may impact response to NUAK inhibition^9^. Here, we used MOBSCs from the amyloid mutant APP/PS1 mice, alongside wildtype littermates, to model firstly, whether we see changes to p-tau Ser356 expression in this AD model, and then the implications of targeting NUAK activity, using WZ4003, under physiological (wildtype) or amyloid mutant (APP/PS1) conditions. Whilst previous work has found that MOBSCs can show accelerated pathological changes compared to *in vivo*^31,32,35,45^, and APP/PS1 mice are found to show increased tau phosphorylation with age^46^, we did not find any differences between the genotypes in this study up to 4 weeks in culture. Nevertheless, the APP/PS1 cultures served an important purpose to establish whether pre-clinical amyloid pathology, or conditions of elevated Aβ production, alter biological responses to WZ4003 treatment. In our work, the response of APP/PS1 MOBSCs was not statistically different to their WT littermates.

A unique aspect of this study is the use of both MOBSCs and HBSCs to examine the impact of WZ4003 treatment. In MOBSCs we found that, whilst WZ4003 treatment lowered both total tau and p-tau Ser356 protein, this reduction coincided with a loss of PSD95, and was proportional to a loss of the neuronal tubulin marker Tuj1. Interestingly, the effects of WZ4003 treatment on tau, synaptic and neuronal protein levels was strongest in the 0-2 week culture period, indicating the early stage cultures were especially sensitive to negative impacts of NUAK inhibition, loss of tau protein, or any potential off-target effects of WZ4003. This possibly reflects differential involvement of NUAK1/2 in different culture phases, or altered processing of tau, over time in culture. By contrast, WZ4003 treatment in HBSCs resulted in a specific reduction in p-tau Ser356, whilst preserving total tau, that occurred alongside *increased* levels of neuronal and synaptic protein. These findings could demonstrate important species differences in how NUAK1 regulates tau and highlight the benefits of using human experimental systems to assess impacts of pharmacological agents^39,47,48^. However, another key difference between MOBSCs and HBSCs is the age of the brain tissue used to generate slices. MOBSCs are taken from postnatal (P6-P9) animals, whilst the age of human brain in this study ranged from 37-76 years old. Therefore, another interpretation of the different response to WZ4003 in MOBSCs versus HBSCs is potential differences in the role of NUAK1/2 during development versus ageing^49^. Indeed, a number of studies have identified key roles of NUAK1 in regulating a number of developmental processes including; axon elongation^44,50,51^, axon branching^50^, and cortical development^51^. It may be that the postnatal slice cultures are negatively affected by either NUAK1/2 inhibition or the loss of total tau during this period, whilst adult human tissue is less dependent on NUAK activity, (or benefits from the relative preservation of total tau levels-which may be important for physiological function^3^). Indeed, our results here indicate NUAK1 inhibition may increase neuronal and synaptic protein levels in adult human brain tissue. One limitation of the present study is that it uses a single small molecule tool which has activity at both NUAK1 and NUAK2, and whilst the published kinase-selectivity data suggests it is a relatively clean inhibitor^30^ we cannot rule out activities at other kinases. Future work should explore further compounds with selectivity for NUAK1 over NUAK2 and more complete profiles.

Human slice cultures represent a translationally powerful new tool for neuroscience research^37–40,47,48,52–56^. Although live human tissue has historically been difficult to obtain, with close collaboration with neurosurgery units, research nurse teams and the laboratory scientists, we have established an efficient pipeline to obtain and culture human brain slices. We show here that they can be an effective tool to examine the impact of pharmacological compounds in live human brain tissue. Previous work has shown benefits of using human cerebrospinal fluid to boost longevity of HBSCs, particularly in regards to electrophysiological activity^37,39^. In our work, using an enriched stem-cell like medium^38^, we find HBSCs retain MAP2 positive neuronal cell bodies and intact neurites, and we are readily able to detect tau, neuronal and synaptic proteins via Western blot for at least 2 weeks *in vitro*. By comparing control and treatment conditions in slices taken from the same individual, we are able to detect biologically relevant responses, even on a background of unavoidable variations in patient age, sex, brain region taken and variations in patient lifestyle and genetic factors. It is worthy of comment that all of our HBSCs, despite none being clinically diagnosed with AD, had detectable levels of p-tau Ser356 in protein extracts, in contrast to our post-mortem study which found very little p-tau Ser356 in Braak 0-I control protein extracts. This could represent that p-tau Ser356 is susceptible to degradation in the post-mortem interval, and thus small levels of p-tau Ser 356 in control post-mortem brain in our samples was rendered undetectable. Alternatively, this could represent live human neurons responding to the culture system itself, such as upregulation of p-tau in response to injury caused during slice culture generation, similar to upregulation of p-tau seen after traumatic brain injury^3,57^. Such differences will be important to reflect on as the tool becomes more widely used. The use of HBSCs as a research tool is expanding, and comparison between mouse tissue, primary cultures, post-mortem human and live human tissue models is likely to be highly valuable when assessing the translational viability of future therapies under development.

In summary, the work in this study further highlights p-tau Ser356 as a potential target of interest in developing AD therapeutics, with increased p-tau Ser356 strongly correlating with Braak stage, being a near-ubiquitous presence in NFTs and co-localising with synapses in AD brain. Whilst NUAK inhibition via WZ4003 treatment of postnatal MOBSCs results in tau lowering that is proportional to loss of synaptic and neuronal protein, we find WZ4003 effectively and specifically lowers p-tau Ser356 in live, adult human brain tissue, highlighting the importance of using complementary experimental systems in pre-clinical work. Future work should further explore the impact of pharmacological NUAK inhibition *in vivo* using a range of tau and amyloid pathology models and with NUAK1 inhibitors optimised for increased potency and drug-like properties with more fully characterised selectivity profiles.

### Availability of Data and Materials

Data is deposited on the University of Edinburgh DataShare digital repository or is available at reasonable request from the corresponding author.

### Author’s Contributions

CD & LT designed the experiments with valuable input from TS-J, JC, JS, SB and PB. LT, ES, CP, OL-S, RM, SM, JR, MS-J, JC, TS-J & CD performed experiments, collected data or performed statistical analysis. HK provided valuable training and advice on the human brain slice culture method. PB & IL provided surgical human tissue samples. CD, PB and SB developed human brain tissue collection and culturing pipelines in Edinburgh. LT and CD wrote the manuscript, and all authors contributed to editing the manuscript and provided feedback. All authors approved the final version of this manuscript.

## Acknowledgements

We would like to thank the University of Edinburgh Bioresearch and Veterinary Services staff for their involvement in breeding and maintaining the mice used in this study. We thank the NRS BioResource and Tissue Governance unit and EMERGE Research Nurse team for their assistance in obtaining informed consent and surplus cortical tissue samples from NHS patients. We thank the MRC Edinburgh Brain Bank for providing human post-mortem tissue. Finally we would like to thank patients and their families for their tissue donations.

## Supplementary Figures

**Supplementary figure 1:**
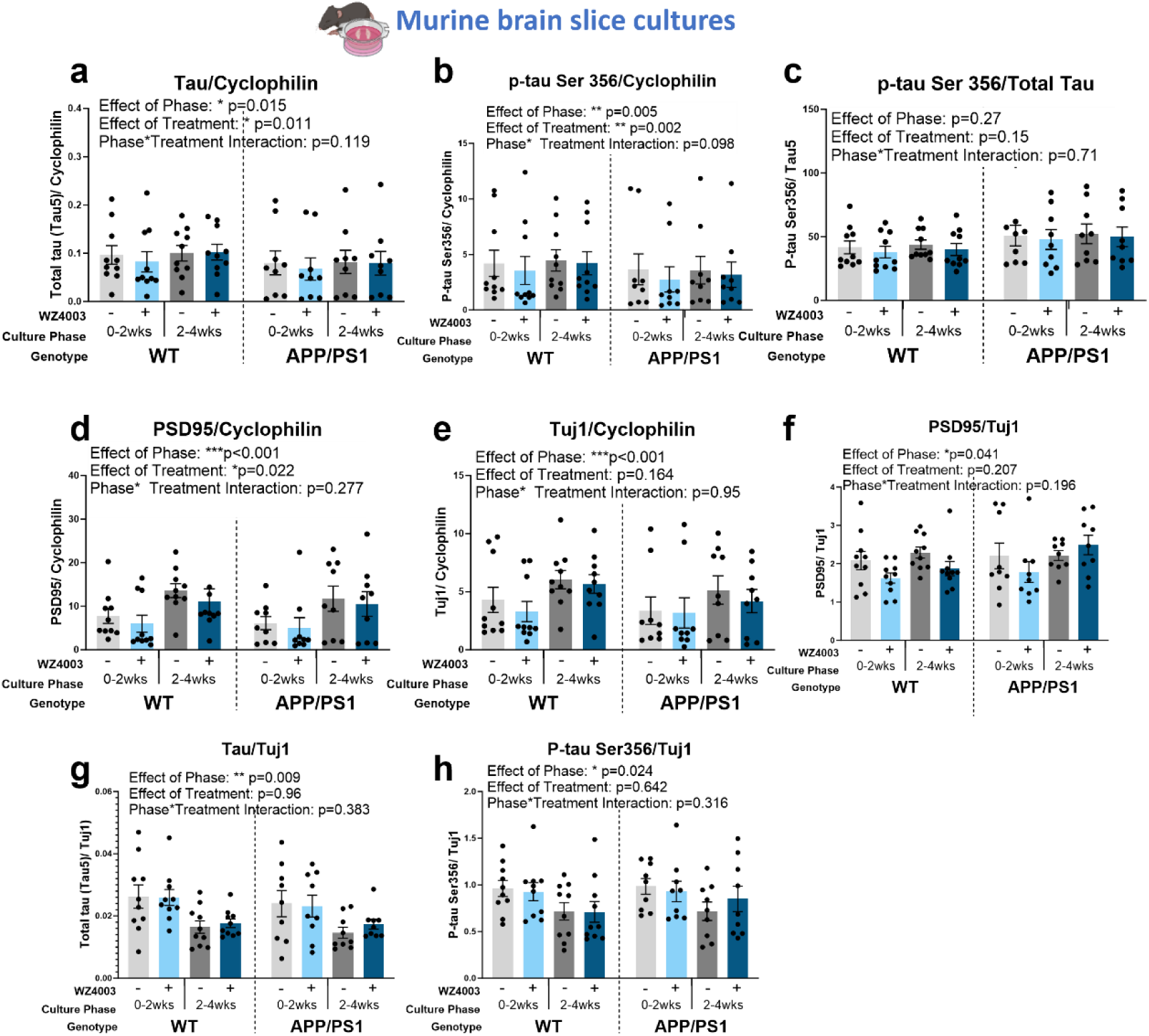
Absolute data graph display for mouse brain slice culture Western blots. *Data and statistics are the same as in Figure 5 and 6, but graphically displayed to show absolute values across mice, not normalised to 0-2 week control to highlight the lack of genotype effects. **(a)** There is a significant effect of phase (*F_(1,51.00)_=6.28, p=0.015) and treatment (*F_(1,51.00)_=7.01, p=0.011) on the levels of total tau, but no effect of genotype (F_(1,21.30)_=0.02, p=0.894). **(b)** There is a significant effect of phase (**F_(1,51.00)_=8.73, p=0.005) and treatment (**F_(1,51.00)_=10.67, p=0.002) on the levels of p-tau Ser356, and a trend interaction between Phase*Treatment (F_(1,51.00)_=2.83, p=0.098), but there is no effect of genotype (F_(1,26.41)_=0.00, p=0.997). **(c)** There are no significant effects of phase (F_(1,51.00)_=1.25, p=0.27), treatment (F_(1,51.00)_=2.10, p=0.15) or genotype (F_(1,26.51)_=0.08, p=0.781) on the ps356/total tau ratio. **(d)** There is a significant effect of phase (***F_(1,51.00)_=26.65, p<0.001) and treatment (*F_(1,51.00)_=5.60, p=0.022) on the levels of PSD95 normalised to cyclophilin, but no effect of genotype (F_(1,41.92)_=0.21, p=0.652). **(e)** There is a significant effect of phase (***F_(1,51.00)_=13.36, p<0.001) on the levels of Tuj1 normalised to cyclophilin, but no effects of treatment (F_(1,51.00)_=1.99, p=0.164) or genotype (F_(1,41.92)_=0.00, p=0.957). **(f)** There is a significant effect of phase (*F_(1,51.00)_=4.39, p=0.041) but no effect of treatment (F_(1,51.00)_=1.63, p=0.207) or genotype (F_(1,47.89)_=0.13, p=0.715) on the levels of PSD95 when normalised to Tuj1. **(g)** There is a significant effect of phase (**F_(1,51.00)_=7.38, p=0.009), but no effect of treatment (F_(1,51.00)_=0.00, p=0.964) or genotype (F_(1,42.38)_=0.08, p=0.774), on the levels of total tau when normalised to Tuj1. **(h)** There is a significant effect of phase (*F_(1,51.00)_=5.40, p=0.024), but no effect of treatment (F_(1,51.00)_=0.22, p=0.642) or genotype (F_(1,42.83)_=0.00, p=0.957), on the levels of p-tau Ser356 when normalised to Tuj1. N=9 APP/PS1 and 10 WT animals, 1-2 slices per animal per condition. Cartoons generated using BioRender.*

**Supplementary figure 2:**
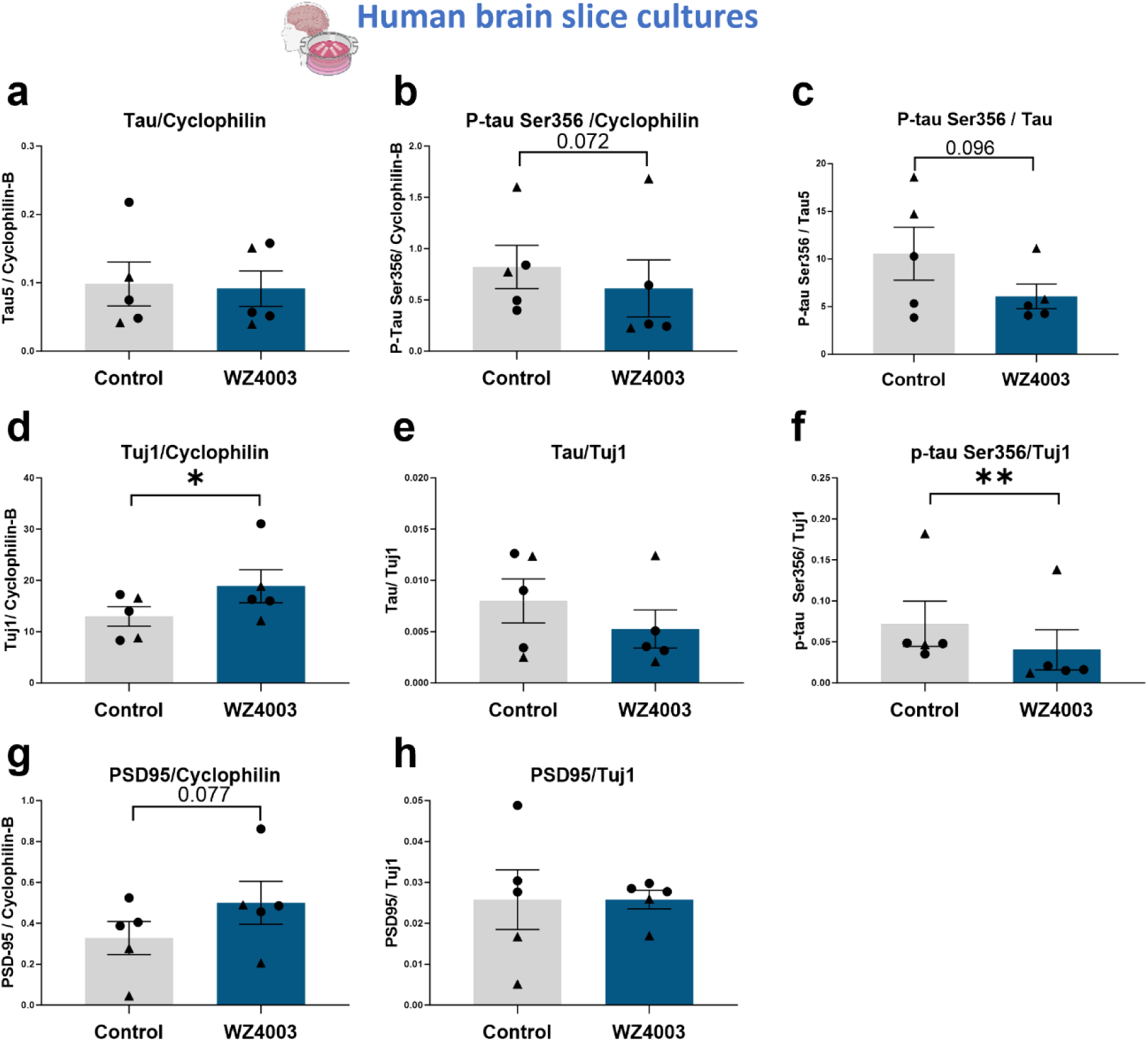
Absolute data graph display for human brain slice culture Western blots. *Data and statistics are the same as in Figure 7, but graphically displayed to show absolute values across cases, not normalised to control. **(a)** WZ4003 does not significantly alter levels of tau (normalised to cyclophilin) (T_(4)_=0.43, p=0.688). **(b)** There is a trend for WZ4003 treatment to reduce p-tau Ser356 (normalised to cyclophilin) (T_(4)_=2.43, p=0.072). **(c)** There is a trend for WZ4003 to reduce the ratio of p-tau Ser356/ total tau (T_(4)_=2.17, p=0.096). **(d)** WZ4003 treatment significantly increased Tuj1 levels (*T_(4)_=3.39, p=0.028). **(e)** WZ4003 does not significantly alter tau levels as a proportion of neuronal protein (T_(4)_=2.04, p=0.11). **(f)** WZ4003 significantly lowers p-tau Ser356 as a proportion of neuronal protein (**T_(4)_=4.81, p=0.0086). **(g)** There is a trend for WZ4003 to increase PSD95 protein (normalised to cyclophilin) (T_(4)_=2.37, p=0.077). **(h)** There is no effect of WZ4003 on PSD95 protein (normalised to Tuj1) (T_(4)_=0.74, p=0.501). N=5 cases per condition. Each point represents an individual human case, triangles= males, circles= females. Cartoons generated using BioRender.*

